# Analysis of molecular and cellular bases of honey bee mushroom body development

**DOI:** 10.1101/2025.02.10.637583

**Authors:** Shuichi Kamata, Takeo Kubo, Hiroki Kohno

## Abstract

Honey bee mushroom bodies (MBs) comprise three class I Kenyon cell (KC) subtypes (lKC, mKC, and sKC) with distinct somata sizes and locations and gene expression profiles. While these KC subtypes have been suggested to function in different behavioral regulations, the molecular and cellular basis of their development remains obscure. Here, we showed that lKCs, mKCs, and sKCs are produced in that order at different pupal stages by labeling proliferating MB cells with 5-ethynil-2’-deoxyuridine at various pupal stages. RNA-sequencing analysis of FACS-sorted pupal MB cells identified genes that were upregulated in proliferating and non-proliferating MB cells, respectively. Furthermore, *in situ* hybridization of some of these genes labeled the proliferating cells or young KCs in the pupal MBs. Our findings reveal the basic scheme of the molecular and cellular processes of honey bee MB development and suggest that they are at least partially different from those of *Drosophila* MB development.

## Introduction

The European honey bee (*Apis mellifera*) is a eusocial insect, and workers exhibit advanced social behaviors, such as age-dependent division of labor and dance communication [1–3]. Honey bees also possess excellent cognitive abilities, such as memorizing flower patches several kilometers away from their hive and learning concepts [2,4,5]. Mushroom bodies (MBs), a higher-order center of the insect brain, have long been studied in honey bees to investigate the molecular and neural mechanisms underlying honey bee social behaviors and cognitive abilities [6–11]. In insects, the MBs are involved in learning, memory, and sensory integration [12,13]. Functional impairment of MBs causes defects in learning and memory in some insects, including honey bees [14–16]. Honey bee MBs change the volume of the neuropil as well as the gene expression profiles associated with the division of labor in workers from nursing their brood inside the hives to foraging for nectar and pollen outside the hives [17–19]. In addition, the MBs of Aculeata, the most recently branching hymenopteran lineage including honey bees, ants, and hornets, are more elaborate in terms of morphology and cytoarchitecture than those of Symphyta, the most basal-solitary hymenopteran lineage [20,21].

Honey bee MBs comprise four types of Kenyon cells (KCs): three subtypes of class I KCs [large-type KCs (lKCs), middle-type KCs (mKCs), and small type-KCs (sKCs)] and class II KCs, which can be distinguished based on the size and location of their somata in the MBs and their gene expression profiles [22,23]. The distinct function of each class I KC subtype was inferred from the function of genes selectively expressed in each KC subtype and the expression patterns of immediate early genes. Genes involved in calcium signaling, such as *Ca2+/calmodulin-dependent protein kinase II* (*CaMKII)* and *1,4,5-inositol trisphosphate receptor* (*IP_3_R*), are preferentially expressed in lKCs [22,24,25], and knockdown of *CaMKII* by RNAi inhibits long-term memory in honey bees, suggesting that lKCs are involved in learning and memory [26]. In contrast, the expression of an immediate early gene, *kakusei*, was upregulated in both mKCs and sKCs during foraging flights, suggesting their role in sensory processing during foraging flights [22,27–29]. Recent studies have suggested that *ecdysone receptor* (*EcR*), which is preferentially expressed in sKCs, is induced in worker brains after foraging flights and regulates the expression of downstream genes related to lipid metabolism [30].

To elucidate the *in vivo* functions of each KC subtype, it is preferable to analyze the behavior of mutants that are defective in MB development and lack a specific subtype. Such mutants have already been generated in *Drosophila*, where the molecular and cellular bases of MB development have been extensively studied. In pupal *Drosophila* MBs, neural stem cells (neuroblasts) divide asymmetrically to produce themselves and ganglion mother cells (GMCs), and GMCs further divide once to produce two KCs [31,32]. While neuroblasts continue to retain the consistent properties to express their marker genes such as *Deadpan* (*dpn*) and *asense* [33], they produce three types of KCs that constitute the *Drosophila* MBs in the order of γ, α’β’, and αβ KCs by changing the expression patterns of certain transcription factors (TFs) at different metamorphosis stages from the late larval to 90 h after pupation [32,34–37]. Subsequently, MB neuroblasts are eliminated via apoptosis [36,37]. Artificially altering the expression levels of genes involved in the production of these three types of KCs results in the ablation of a specific subtype [33].

In the honey bee, MB neuroblasts are located in the inner core of the MB calyces and sequentially produce class I KCs from the outer to the inner cells in the adult MBs until the mid-pupal stages [38]. Expression analysis of some marker genes of lKCs revealed that lKCs were produced the earliest among the three KC subtypes at the early pupal stages [39]. MB neuroblasts disappear due to apoptosis at the mid-pupal stage on the 5th day after pupation (P5), and KC production ceases in subsequent pupal stages [40]. Recently, a honey bee-specific non-coding RNA, *Nb-1*, was reported to be preferentially expressed in the proliferating cells of MBs [41]. However, little is known about the molecular and cellular basis of the honey bee MB development, even whether they share some mechanisms with the *Drosophila* development. This makes it difficult to identify genes involved in the production and development of each KC subtype for the future production of mutant honey bees lacking a specific KC subtype, although methods for CRISPR/Cas9 mutagenesis have been established in the honey bee.

In the present study, we aimed to clarify when each honey bee KC subtype is produced during metamorphosis and to analyze which genes, especially TFs, are related to the production and maturation of KCs in honey bees. We identified the pupal stages at which each subtype is produced by labeling proliferating cells in the pupal MBs with 5-ethynil-2’-deoxyuridine (EdU) at various pupal stages and subsequent detection of EdU in the brain after emergence as adults. We also identified genes differentially expressed in both proliferating and non-proliferating cells in the pupal MBs using RNA-sequencing (RNA-seq) analysis. The expression pattern analysis of some genes indicated the distribution of neuroblasts, immature KCs, and mature KCs in the MBs at various pupal stages. Our findings provide a basic scheme for the molecular and cellular processes of class I KC development in the honey bee MB, suggesting that these processes are at least partially different from those in *Drosophila*.

## Results

### Identification of pupal stages in which each class I KC subtype is produced

So far, honey bee pupal stages have been classified as P1 to P9 (each approximately 1 d long) based on the coloring of compound eyes and bodies [40,42,43]. However, this classification is rather subjective and inaccurate and has insufficient temporal resolution, considering the possibility that a certain KC subtype is produced in a period shorter than one day. Therefore, we constructed a new experimental system to artificially rear the last instar larvae collected from bee hives on a 24-well plate in a styrofoam box with controlled internal temperature (34L) and relative humidity (70%), while taking a picture of them every 10 min with an infrared camera to identify the pupation time (**Figure S1**). The identified pupation time allowed us to determine the exact pupal stage, that is, the time after pupation, of each individual with a 10-min temporal resolution.

Using the pupae whose exact pupal stages were identified in this system, we investigated when each KC subtype was produced (**Figure 1A**). Proliferating MB cells were labeled by injecting EdU into the heads of pupae at various stages up to 120 h after pupation (hap), and the pupae were reared in an incubator (34℃) till emergence. Using the brain sections of the adults just after emergence, both fluorescence *in situ* hybridization (FISH) for *juvenile hormone diol kinase (jhdk*), a gene whose expression is strong in lKCs, moderate in sKCs, and scarce in mKCs [44], and EdU detection were performed to examine the KC subtypes in which EdU signals were detected (**Figures 1B-D and S2**). In all individuals, *jhdk* signals were detected in both the peripheral and inner core areas inside the MB calyx, which corresponded to lKCs and sKCs, respectively, but not in the area between them, which corresponded to mKCs [44]. EdU signals were detected in the peripheral area inside the MB calyx of pupae injected with EdU at 7 or 14 hap, which corresponded to lKCs in the merged image of EdU and *jhdk* signals, suggesting that MB neuroblasts produced cells that differentiated into lKCs during these pupal stages (**Figures 1B, 1E and S2A**). EdU signals were detected in areas between the peripheral and inner core areas inside the MB calyx of pupae injected with EdU at 30, 32, 42, and 49 hap, which corresponded to mKCs (30 and 32 hap) and mKCs and sKCs (42 and 49 hap), respectively, suggesting that neuroblasts produced cells that differentiated into these KC subtypes during these pupal stages (**Figures 1C, 1E and S2B-D**). EdU signals were detected in the inner core area inside the MB calyx of pupae injected with EdU at 85, 88, 90, and 108 hap, which corresponded to sKCs, suggesting that neuroblasts produced cells that eventually differentiated into sKCs during these pupal stages (**Figures 1D, 1E and S2E-G**). These results strongly suggest that lKC, mKC, and sKC were produced in that order during the pupal stage (Figure 1E). In contrast, in some individuals, EdU signals were detected in multiple subtypes (42 and 49 hap) (**Figure 1E and S2C-D**). Because the injected EdU is thought to remain in the brain for several hours, it is plausible that EdU was injected into these individuals during periods when the KC subtypes to be produced were just switching. Therefore, we concluded that lKCs, mKCs, and sKCs are produced, in this order, at different pupal stages: until approximately 30 hap (the pupal stage when lKCs are produced; P-lKC), between 30 and 50 hap (P-mKC), and between 50 and 120 hap (P-sKC).

**Figure 1.**
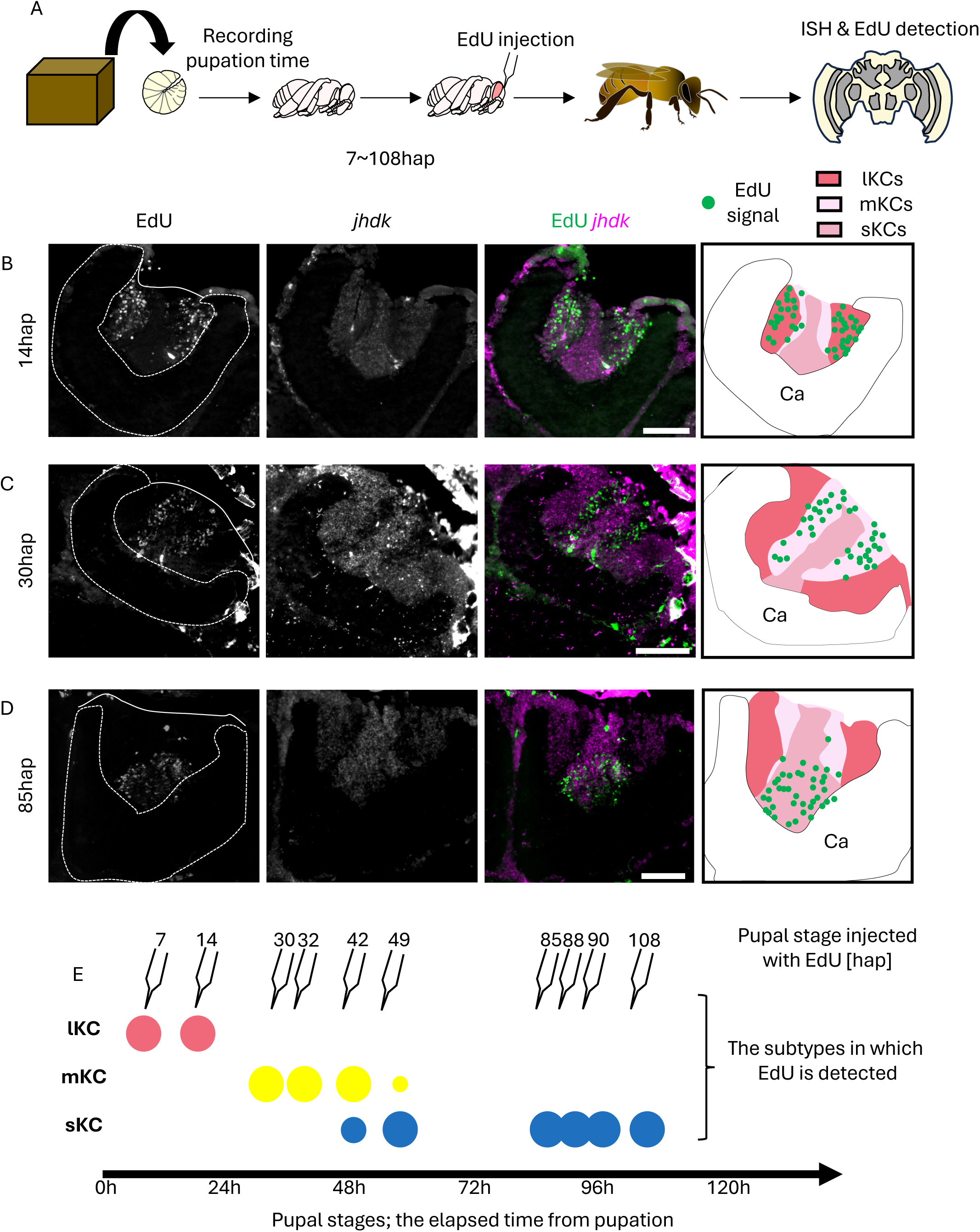
Identification of the pupal stage when each subtype is produced from neuroblasts. (A) A scheme of the experimental design. Last instar larvae were collected from the bee hives and EdU was injected into the pupal heads at various pupal stages. After emergence, EdU detection and ISH in the brain sections were performed. (B-D) Distribution of EdU signals in the brain of adult individuals injected with EdU at 14hap (B), 30hap (C), and 85hap (D). From left to right panels, the EdU signals, *jhdk* FISH signals, merged signals, and schematic diagrams of EdU signals and each subtype are shown. Ca, a calyx in the MB. Scale bar; 100 μm. (E) Correspondence between the pupal stage injected with EdU and the subtypes where EdU signals were detected. The size of each circle represents the approximate proportion of EdU-detected cells in each subtype.

### Investigation of gene expression profiles of proliferating and non-proliferating pupal MB cells by RNA-seq analysis

To elucidate the molecular basis of honey bee MB development, it is important to identify the genes expressed in each cell type constituting the developing MBs. Therefore, we searched for genes preferentially expressed in proliferating cells (neuroblasts and GMCs) and non-proliferating cells (KCs and glia) in pupal MBs. We used pupae at the P-lKC in this experiment because they can be prepared earlier than those at the P-mKCs and P-sKCs. MB cells dissected from pupae were dispersed and stained with Hoechst33342 for DNA staining, and cell cycle analysis was performed using fluorescence-activated cell sorting (FACS) (**Figure 2A**). Two peaks were detected in the histogram of nuclear staining intensities, and the minor peak had a DNA content approximately twice that of the major peak (**Figure 2B**). The major and minor peaks were considered to correspond to cells with nuclear phases of 2X and 4X, respectively. We performed RNA-seq analysis using the isolated 2X (average of approximately 30,500 cells in three lots) and 4X (average of approximately 5,500 cells in three lots) fractions and identified 148 and 204 differentially expressed genes (DEGs, FDR < 0.05, and over two-fold change) that were upregulated either in 2X or 4X fraction, respectively (**Table S1**). To validate the result of cell cycle analysis, we performed GO enrichment analysis using these DEGs and found that DEGs upregulated in 2X fraction were enriched with genes related to neuronal maturation processes, such as “axonogenesis” (ranked 1st) and “synapse organization” (ranked 3rd) (**Figure 2C**). In contrast, DEGs upregulated in the 4X fraction were enriched with genes related to some neurogenesis processes, such as “cell fate commitment” (ranked 2nd), “generation of neuron” (ranked 4th), and “mitotic centrosome separation” (ranked 6th) (**Figure 2D**). Therefore, we concluded that non-proliferating or proliferating cells were successfully enriched in the 2X or 4X fraction, respectively, and comprehensive gene expression profiles of these cells were obtained.

**Figure 2.**
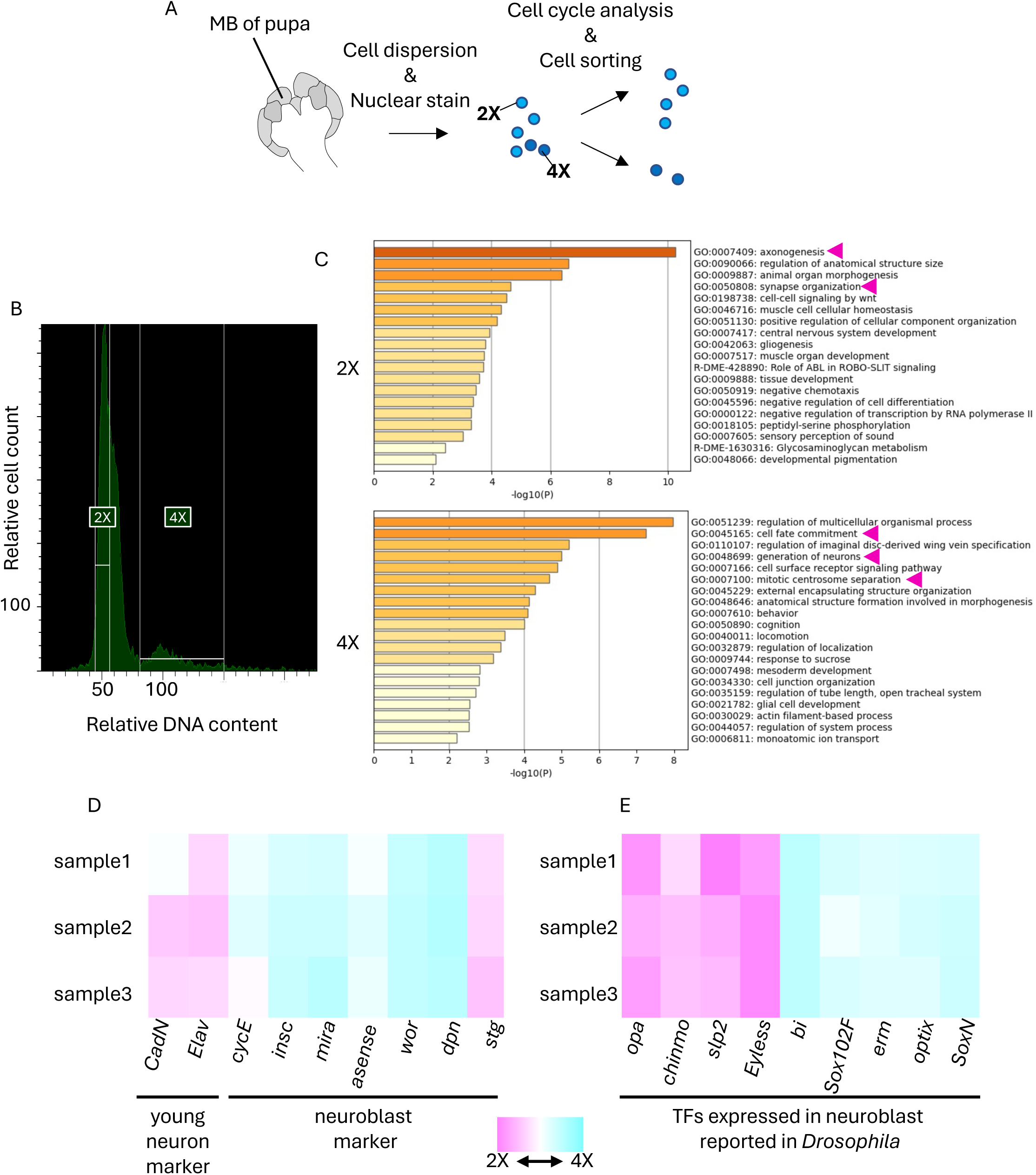
RNA-seq analysis of 2X and 4X cell fractions in the pupa MBs sorted by FACS. (A) A scheme of experimental design. MBs were dissected from pupae at P-lKCs. After cell dispersion and nuclear staining with Hoechst33342, both the 2X and 4X fractions were sorted using FACS. (B) Cell cycle analysis of dispersed pupal MB cells. The ranges of relative DNA content (the intensity of Hoechst33342) of cells sorted as the 2X or 4X fraction are indicated with white lines. (C) The results of GO enrichment analysis using DEGs in each fraction by Metascape. Magenta arrows indicate the GO terms mentioned in the text. (D, E) Heatmap showing the relative expression levels of marker genes of young neurons or neuroblasts (D) and genes coding transcription factors (TFs) reported to be expressed in neuroblasts in *Drosophila* (E). The relative values are calculated for each sample by comparing the expression levels of genes in the 2X and 4X fractions. Magenta and blue show that expression levels are higher in the 2X and 4X fractions, respectively.

Previous studies in *Drosophila* identified marker genes for neuroblasts [*string* (*stg*), *deadpan* (*dpn*), *worniu* (*wor*), *asense*, *miranda* (*mira*), *inscrutable* (*insc*), *cyclinE* (*CycE*)] and young neurons [*embryonic lethal abnormal vision* (*elav*), *Cadherin-N* (*CadN*)] in the development of the central nervous system [45–48]. When the expression levels of honey bee orthologs of these genes were compared between the 2X and 4X fractions, we found that the expression levels of the marker genes for neuroblasts and young neurons, with the exception of *stg*, tended to be high in the 4X and 2X fractions, respectively (**Figure 2D**). In addition, *wor*, *mira*, and *insc* were included in the DEGs upregulated in the 4X fraction, whereas *stg*, *elav*, and *CadN* were included in those in the 2X fraction. In mammals and *Drosophila*, a variety of TFs are sequentially expressed in neural cells from the time the cell is produced until maturation, leading to the acquisition of adult gene expression profiles [49–51]. Therefore, we focused on some TFs that are expressed in neuroblasts during development in *Drosophila* and examined their expression levels in the 2X and 4X fractions. We found that *SoxNeuro* (*SoxN*) [52]*, Optix* [53], *wor* [54]*, earmuff* (*erm*) [55], and *Sox102F* [56] were included in DEGs upregulated in the 4X fraction, while *Eyeless* [57]*, sloppy-paired* (*slp2*) [58]*, chronologically inappropriate morphogenesis* (*chinmo*) [34], and *odd-paired* (*opa*) [59] were included in DEGs upregulated in the 2X fraction (**Figure 2E**). These results suggest that there are both conserved and unique gene expression profiles in the neuroblasts of honey bees compared to those in *Drosophila*.

### *In situ* hybridization (ISH) of genes with conserved / unique expression patterns in the honey bee pupal MBs

To further investigate the molecular and cellular basis of honey bee MB development, we performed ISH of the genes for some TFs whose expression patterns were suggested to be common with or different from *Drosophila* by RNA-seq, using honey bee pupal brains.

First, we focused on genes suggested to be expressed in the proliferating cells of the honey bee, similar to *Drosophila*. We selected *SoxN* and *optix* as the genes for TFs included in DEGs upregulated in the 4X fraction (**Figure 2E**). Although not included in the DEGs, we also selected *asense* among the known MB neuroblast marker genes of *Drosophila* (**Figure 2D**) because *asense* encodes a TF and is preferentially expressed in MB neuroblasts in *Drosophila* brain [33]. Although *dpn* encodes a TF that is preferentially expressed in *Drosophila* MB neuroblasts [33], such as *asense*, we excluded it from the genes for ISH because its expression level in the honey bee pupal brains was considered too low to be detected by ISH, based on the results of RNA-seq analysis (**Table S1**). ISH of these genes was conducted using brain sections of pupae at four distinct pupal stages: P-lKC, P-mKC, P-sKC, and P7 (based on the coloring of the body and compound eyes according to previous criteria) as a control stage when cell proliferation in MBs was terminated [40]. Signals of *SoxN, optix,* and *asense* were detected at the center of each calyx in all pupal stages except P7, where no signals were detected (**Figures 3A-C and S3A-C**). Weak signals of *asense* were also detected in the area surrounding the inner core (**Figure 3C**). Next, we examined whether these genes were expressed in proliferating cells by detecting ISH and EdU signals in the brain sections of pupae with proliferating MB cells (around P-mKC) injected with EdU 2 h prior to brain dissection. The expression patterns of *SoxN* and *optix* mostly overlapped with the distribution of the EdU signals (**Figure 3D, E**). As for *asense*, the areas with stronger signals, but not those with weaker signals, mostly overlapped with those with EdU signals (**Figure 3F**). These results indicate that *SoxN*, *optix*, and *asense* are mainly expressed in proliferating MB cells during the pupal stages, when all class I KC subtypes are produced, but not in non-proliferating cells (though weakly so for *asense*).

**Figure 3.**
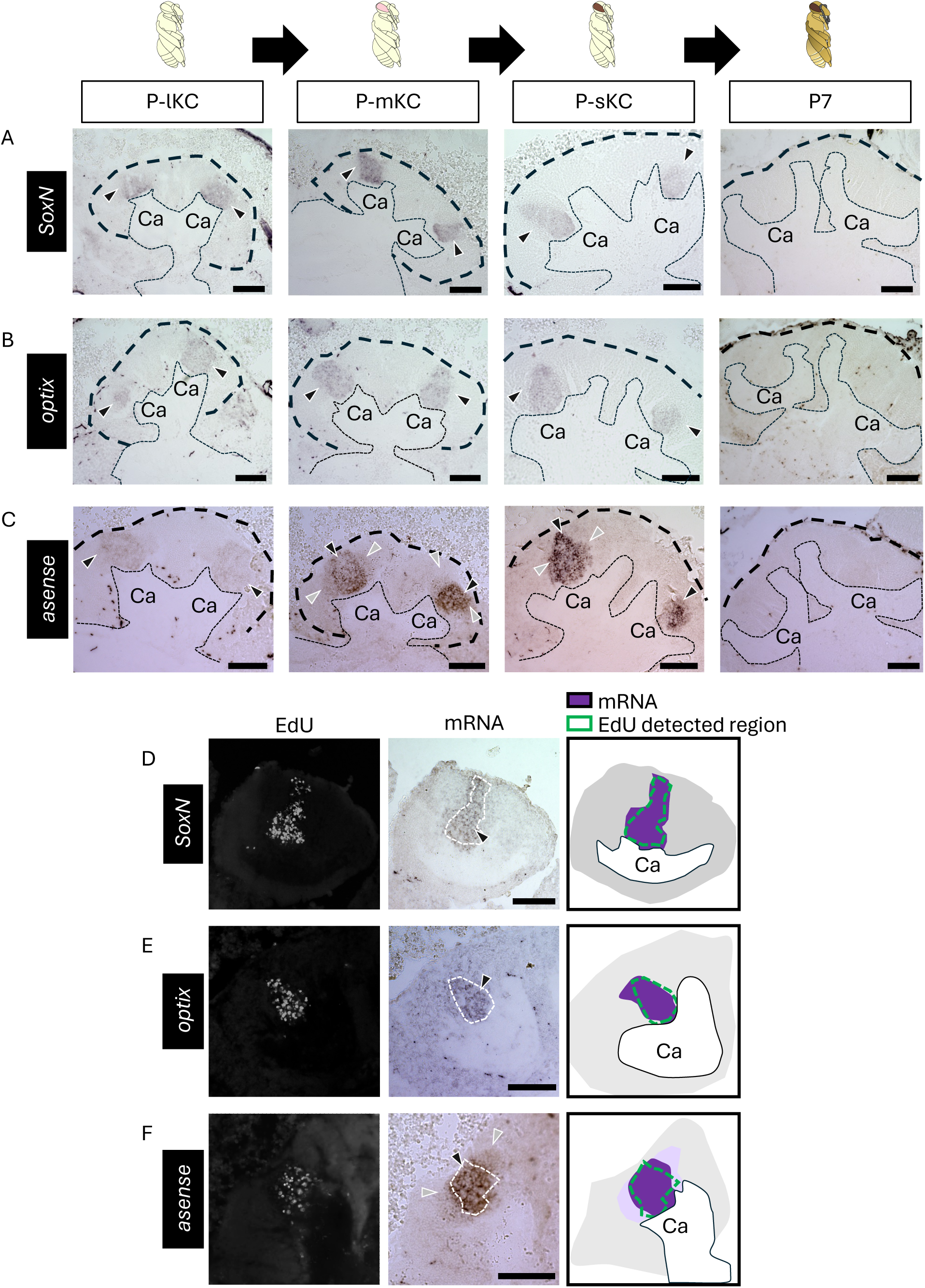
ISH of the genes suggested to be expressed in the proliferating cells in the pupal MBs. (A-C) The results of ISH of *SoxN* (A), *optix* (B), and *asense* (C) in the MBs of pupae at P-lKC, P-mKC, P-sKC, and P7 pupal stage, respectively. The thin and bold black dashed lines indicate the calyces and areas of KC somata, respectively. (D-E) The results of EdU detection and ISH of *SoxN* (D), *optix* (E), and *asense* (F) using the pupae with MB proliferating cells injected with EdU 2 h before dissection. From left to right panels, the EdU signals, ISH signals, and schematic diagrams of EdU and ISH signals are shown. The white dashed lines in ISH images indicate the region where EdU signals were detected. The intensity of the purple color in schematic diagrams represents the differences in the intensity of ISH signals. Black and gray arrowheads indicate strong and weak ISH signals, respectively. Ca, a calyx in the MB. Scale bar; 100 μm.

Next, we focused on genes suggested to be expressed in non-proliferating cells in honey bee MBs, which are inconsistent with the expression patterns known in *Drosophila*. We performed ISH of *opa* using brain sections from pupae at four pupal stages: P-lKC, P-mKC, P-sKC, and P7. The signals of *opa* were weakly detected at the center of each calyx and were strongly detected in the area surrounding the inner core area in all pupal stages except P7, where no signals were detected (**Figures 4A and S3D**). In the pupae (around P-mKCs), whose proliferating cells were labeled with EdU, the area with weak signals of *opa* mostly overlapped with the distribution of cells with EdU signals, but those with strong signals did not (**Figure 4B**). Considering that newly born KCs are located in the area surrounding proliferating cells and are gradually pushed to the outer area inside the calyx by those produced later [34], and that ISH signals of *opa* were not detected at P7, it is suggested that *opa* is preferentially expressed in immature KCs but not in mature KCs.

**Figure 4.**
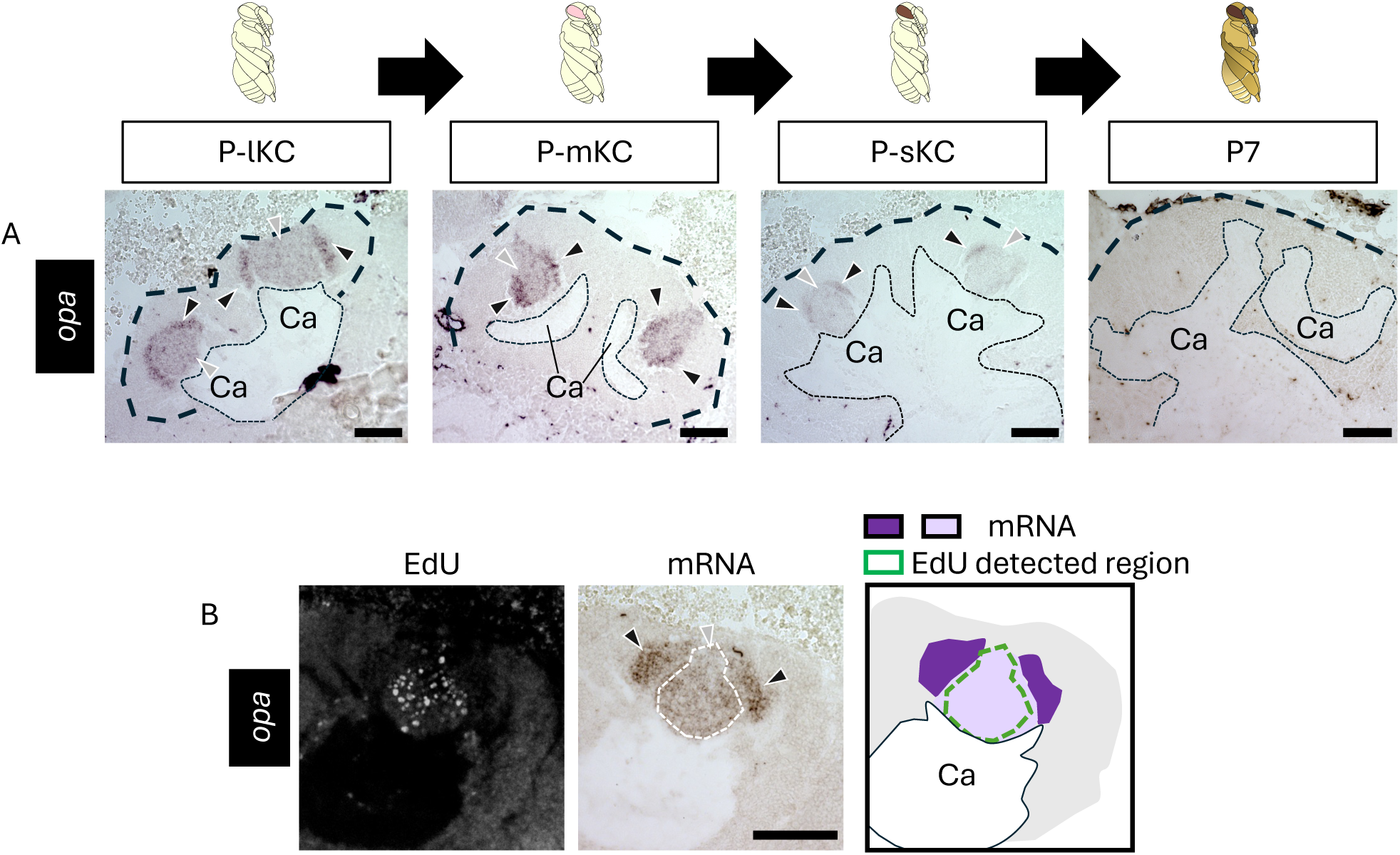
ISH of *opa* in the pupal MBs. (A) The results of ISH of *opa* in the MBs of pupae at P-lKC, P-mKC, P-sKC, and P7 pupal stage, respectively. The thin and bold black dashed lines indicate the calyces and areas of KC somata, respectively. (B) The results of EdU detection and ISH of *opa* using the pupae with MB proliferating cells injected with EdU 2 h before dissection. From left to right panels, the EdU signals, ISH signals, and the schematic diagram of EdU and ISH signals are shown. The white dashed line in the ISH image indicates the region where EdU signals were detected. The intensity of the purple color in the schematic diagram represents the differences in the intensity of ISH signals. Black and gray arrowheads indicate strong and weak ISH signals, respectively. Ca, a calyx in the MB. Scale bar; 100 μm.

To further test this possibility, the distribution of mature KCs in pupal MBs must be determined. Therefore, we analyzed the expression patterns of genes known to be preferentially expressed in each KC subtype in the adult brain. The distribution of lKCs in pupal MBs has been reported by ISH of *Mblk-1*, a marker gene for lKCs, which is expressed strongly in mature KCs and weakly in proliferating cells at P3 [39]; however, the distribution of mKCs and sKCs in pupal MBs has not yet been reported. Therefore, we focused on *MOXD1* and *Frq1*, which are marker genes for mKCs and sKCs, respectively [60], and performed ISH of these genes using brain sections of pupae at four pupal stages: P-lKC, P-mKC, P-sKC, and P7 (**Figure 5A, B**). The signals for *MOXD1* were detected at the periphery, but not in the inner core area inside each calyx at P-mKC and P-sKC, and at the area of mKCs at P7, whereas no signals were detected at P-lKC (**Figure 5A**). In pupae at P-mKC, whose proliferating cells were labeled with EdU, signals of *MOXD1* were detected at the periphery of the EdU signal area (**Figure 5C**). These expression patterns indicate that mature KCs expressing *MOXD1*, i.e., mKCs, were localized in areas distinct from proliferating cells. In contrast, signals of *Frq1* were detected at the center and bottom of each calyx at P-sKC and in the area of sKCs at P7, but not at P-lKC and P-mKC (**Figure 5B**). In the pupae at P-sKC, signals of *Frq1* were detected in the area mostly overlapping with EdU signals (**Figure 5D**), suggesting that *Frq1* begins to be expressed in proliferating cells at P-sKC and continues to be expressed in mature KCs (sKCs). In contrast to *Frq1*, *opa* was not detected at P7 (**Figure 4A**), suggesting that *opa* is transiently expressed during KC maturation.

**Figure 5.**
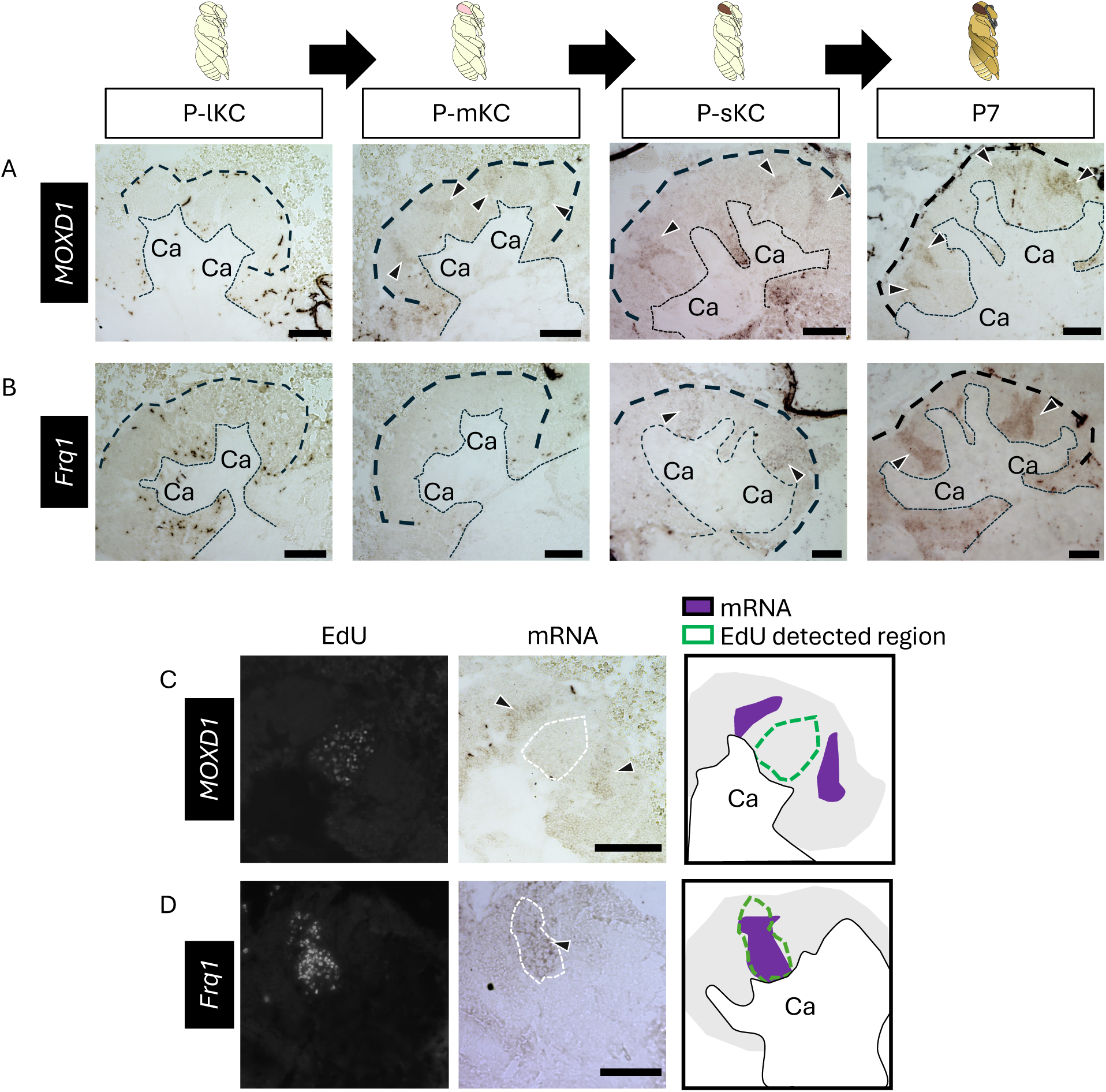
ISH of the genes suggested to be expressed in the mature KCs in the pupal MBs. (A, B) The results of ISH of *MOXD1* (A) and *Frq1* (B) in the MBs of pupae at P-lKC, P-mKC, P-sKC, and P7 pupal stage, respectively. The thin and bold black dashed lines indicate the calyces and region of KC somata, respectively. (C, D) The results of EdU detection and ISH of *MOXD1* (C) and *Frq1* (D) using the pupae at P-mKC (C) and P-sKC (D) injected with EdU 2 h before dissection. From left to right panels, the EdU signals, ISH signals, and schematic diagrams of EdU and ISH signals are shown. The white dashed lines in ISH images indicate the region where EdU signals were detected. Black arrowheads indicate ISH signals. Ca, a calyx in the MB. Scale bar; 100 μm.

Finally, we performed ISH of *opa* and *MOXD1* using serial brain sections of pupae at P-mKC, whose proliferating cells were labeled with EdU, to examine whether *opa* was preferentially expressed in immature KCs. Between the regions where *MOXD1* and EdU signals were detected, there was another population of cells with neither EdU nor *MOXD1* signals (**Figure 6A**). In contrast, a strong *opa* signal was detected in the area adjacent to the area with EdU and weak *opa* signals (**Figure 6B**). Most of the areas where signals of *opa* were detected did not overlap with those of *MOXD1* (**Figure 6C**). These results suggest that cells with strong *opa* signals in the pupal MBs were immature KCs.

**Figure 6.**
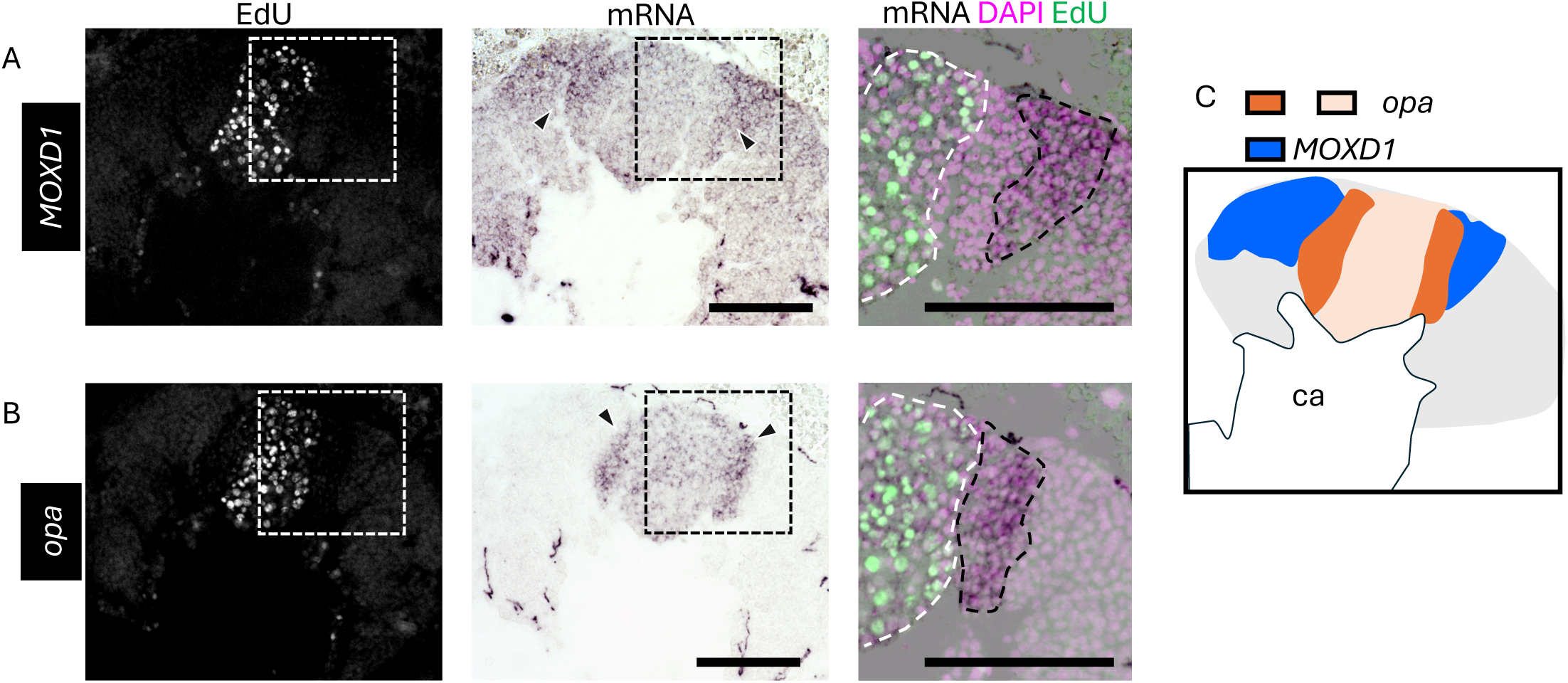
ISH of *opa* and *MOXD1* using serial sections of the pupal MBs. (A, B) The results of ISH of *MOXD1* (A) and *opa* (B) with serial sections of the MBs of pupae at P-mKC injected with EdU 2 h before dissection. From left to right panels, merged images of DAPI and EdU signals, ISH signals, and magnified views of the merged images of DAPI, EdU, and ISH signals in the squares surrounded with white dushed lines are shown. Black arrowheads in the middle panels indicate strong ISH signals. Areas surrounded by black and white dashed lines in the right panels indicate areas with strong ISH signals and those with EdU signals, respectively. (C) The schematic diagrams of the expression pattern of *opa* (dark and light orange) and *MOXD1* (blue). Ca, a calyx in the MB. Scale bar; 100 μm.

## Discussion

In the present study, we identified the exact pupal stage at which each subtype is produced by combining newly established systems for artificial rearing that allow us to identify the time after pupation at a 10-min temporal resolution, and an experiment to label proliferating cells in pupal MBs using EdU. We found that the production of mKCs from neuroblasts occurred only within 24 h, which could not be revealed using the conventional classification of pupal stages (P1-P9) that has a temporal resolution of one day [40,42,43]. We also suggest that by identifying DEGs highly expressed in proliferating and non-proliferating cells, the gene expression profile of honey bee MB-proliferating cells is at least partially different from that in *Drosophila* [33,34,48]. Finally, ISH of some genes in the brains of various pupal stages revealed the distribution and partial gene expression profiles of proliferating cells, immature KCs, and mature KCs in honey bee pupae (summarized in **Figure 7A**). By combining the previous ISH results of *Mblk-1* [39] and *mKast* [23,27], which are preferentially expressed in mature lKCs and mKCs, respectively, and the present ISH results, we propose the differentiation trajectory of KCs from proliferating cells in the honey bee pupal MBs including genes specifically expressed in each cell type (**Figure 7B**): The proliferating cells including neuroblasts and GMCs are located at the inner core area inside the calyces and express *SoxN*, *optix*, and *asense*. The immature KCs produced from proliferating cells are located in the peripheral area adjacent to the area of proliferating cells and express *opa*, but not marker genes, for adult KC subtypes. Mature KCs are then pushed to and located at the most peripheral area and begin to express marker genes for adult KC subtypes, such as *Mblk-1* in lKCs [39], *mKast* and *MOXD1* in mKCs [23,27,60], and *Frq1* in sKCs [60].

**Figure 7.**
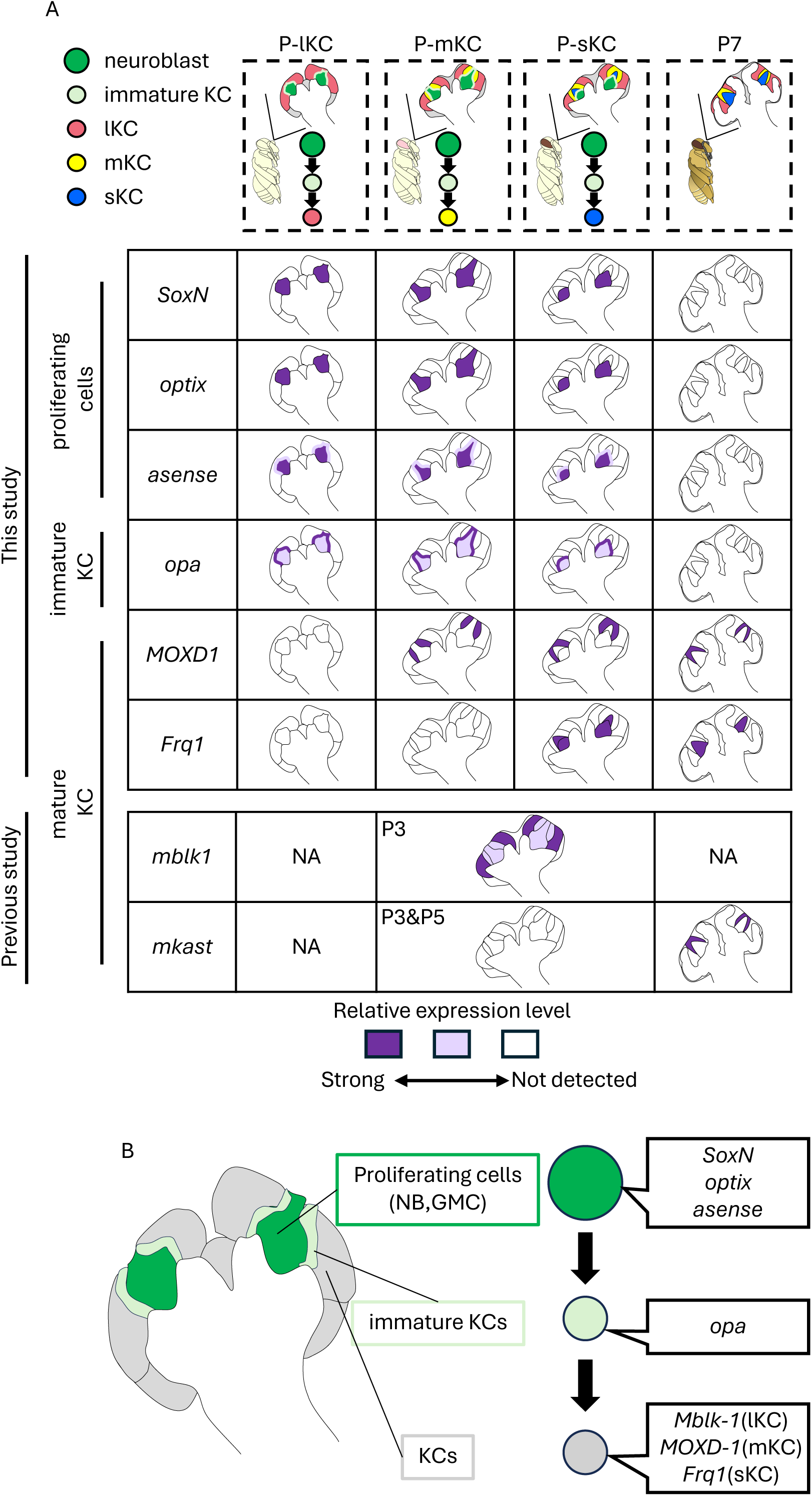
The molecular and cellular developmental basis of class - KCs in the honey bee. (A) Summary of gene expression patterns in the MBs at 4 pupal stages (P-lKC, P-mKC, P-sKC and P7). The intensity of the purple color in each schematic diagram represents the differences in the expression level of each gene. In the previous studies [22,39,66], ISH of *Mblk-1* and *mKast* was performed in pupae at P3 and at P3, P5, and P7, respectively. NA: not analyzed in the previous studies. (B) The differentiation trajectory of KCs from proliferating cells in the honey bee pupal MBs including genes specifically expressed in each cell type.

*SoxN* in *Drosophila* and its vertebrate homolog *Sox-2* in mammal*s* are involved in the maintenance of neural stem cell properties [61–63]. In addition, one of the direct transcriptional targets of *Sox-2* is *SIX3*, the vertebrate homolog of *optix*, which is involved in proper forebrain formation by repressing *Wnt1* signaling in zebrafish [64]. Because *SoxN* and *optix* are expressed in proliferating cells in honey bee pupal MBs, it is possible that *SoxN* also functions in maintaining the stem cell properties of honey bee MB neuroblasts through the regulation of the expression of *optix*. The function of *opa* in optic lobe development has been well-studied in *Drosophila* [48,51,57,65]. Acting within the transcriptional cascade in early- and middle-aged neuroblasts, *opa* promotes the transcription of *Eyeless* and *Oaz*, thereby endowing neuroblasts with temporal characteristics [51]. In honey bee MBs, *opa* is expressed in immature KCs, and both *opa* and *Eyeless* were identified as DEGs that were highly expressed in non-proliferating cells, implying that the transcriptional cascade downstream of *opa* is suppressed in neuroblasts and induced in KCs only transiently before maturation. In addition, considering that *SoxN* expression is repressed by *Eyeless* in *Drosophila* [48], we hypothesized that suppression of *opa* in neuroblasts led to repression of the transcription of *Eyeless* and finally to sustained expression of *SoxN* throughout all pupal stages when class I KCs are produced in the honey bee. Future functional elucidation of these genes will provide important insights into the molecular basis of KC production and maturation in honey bees.

Previous studies have shown that *mKast*, a marker gene for mKCs, is expressed in MBs from P7 [27,66], when KCs production is already complete [40] (**Figure 7A**). Based on this finding, mKCs were previously thought to be originally produced from MB neuroblasts as the same KCs that differentiate into lKCs or sKCs, and subsequently follow a differentiation trajectory to mKCs that is distinct from those to lKCs or sKCs [22,23]. However, the present study revealed that *MOXD1*, another marker gene of mKCs [60], is already expressed in mature KCs at the P-mKC (**Figure 5A**), strongly suggesting that mKCs acquire a unique gene expression profile earlier than previously thought after production from MB neuroblasts. *mKast* may be expressed downstream of an unidentified transcriptional cascade involved in mKC maturation.

In the present study, we identified *SoxN*, *optix*, and *asense* as candidate genes that are commonly involved in the production and maturation of all KC subtypes (**Figure 7B**). However, the TFs responsible for inducing the distinct gene expression profiles in each KC subtype remain unclear. In *Drosophila* KCs, the expression of the TF *Chinmo* in neuroblasts decreases gradually during development, leading to the production of distinct KC types depending on the developmental stage [34]. Therefore, it is plausible that each KC subtype in honey bee MBs is also produced from neuroblasts in a temporally regulated manner. However, there is another possibility that homogeneous immature KCs common to all lKCs, mKCs, and sKCs are produced regardless of the pupal stage, and then differentiate into each KC subtype through distinct cell fate determination processes during maturation. These should be examined in the future by searching for genes that are selectively expressed in MB neuroblasts at the pupal stages of P-lKC, P-mKC, and p-sKC. Nonetheless, it may be possible to create honey bees deficient in each KC subtype by suppressing functions of TFs, which are identified as commonly expressed in NBs at any pupal stages, in a pupal stage-specific manner.

The present study suggests that, although there is a large degree of overlap in the gene expression profiles of proliferating cells between honey bees and *Drosophila*, the expression patterns of some genes, such as *opa* are not consistent between these two species. The differences in brain structures, such as MB size, and behavioral properties between *Drosophila* and honey bees [67,68], may be at least partly due to these differences in the molecular and cellular basis of MB development. Future evo-devo studies on molecular and cellular MB development in several insect species may elucidate the molecular developmental basis of MBs, which may lead to the evolution of a wide variety of behavioral traits.

## Limitations of the study

EdU is a thymidine analog that is incorporated into the genomic DNA of cells during DNA synthesis; therefore, we could not distinguish between neuroblasts and GMCs using EdU signal patterns. The cells of the 4X fraction sorted by FACS were thought to be in the S, G2, and M phases of the cell cycle; therefore, DEGs upregulated in 4X fractions are thought to be expressed in both neuroblasts and GMCs. Further analysis is required to identify genes that can be used as distinct molecular markers of neuroblasts or GMCs.

## Methods

### Animals

European honey bee (*Apis mellifera* L.) colonies were purchased from Rengedo Beekeepers (Saga, Japan) or Kumagaya Beekeepers (Saitama, Japan) and maintained at The University of Tokyo (Tokyo, Japan). For the artificial rearing of the last instar larvae into adult workers, a comb-frame containing the last instar larvae was collected from the hive and placed in an artificial rearing container (wooden box) in the laboratory, kept at 34L for about half a day. The last instar larvae that emerged from cells of comb-frame and fell onto the bottom of the container were transferred onto a 24-well plate with paper towels on the bottom of each well, and reared in a styrofoam box at a temperature of 34℃ and relative humidity of 70%, controlled by a heat panel (Midori Sangyo) and a saturated NaCl solution. An infrared camera (Kumantech) was set at the top of the box, and images were captured every 10 min. After pupation, the individuals were maintained in an incubator (NK System) under the same conditions and subjected to EdU injection, cell dispersion, and gene expression pattern analyses.

### EdU Injection

Under a binocular microscope, the pupal head cuticle near the ocelli was carefully pierced using an insect pin and a glass capillary (Drummond) filled with 10 mM EdU in phosphate-buffered saline (PBS; 136 mM NaCl, 2.68 mM KCl, 10.1 mM Na2HPO4, 1.77 mM KH2PO4) with a tip diameter of approximately 20 μm was then inserted through the hole to a brain in the depth of about 2 mm. After injection of 1 µl of EdU solution, the pupae were kept again in the incubator at 34L and 70% relative humidity for 2 h after injection or until within 2 days after emergence.

### Double staining with EdU detection and fluorescent / chromogenic ISH

After anesthetizing on ice, the brains of pupae or adults were dissected in insect saline (130 mM NaCl, 5 mM KCl, 1 mM CaCl2) using a microsurgical scalpel and tweezers under a binocular microscope, embedded in Tissue-Tek OCT compound (SAKURA Finetek, Japan) without fixation, immediately frozen on dry ice, and stored at −80L until use. ISH was performed as previously described [27]. The brain samples were sliced into thin sections with a thickness of 20 μm for adults and 15 μm for pupa using a cryostat (Leica or Thermo Fisher), attached onto a MAS-coated glass (Matsunami glass), air-dried for at least 1 h, and fixed in 4% paraformaldehyde in phosphate buffer (PB; 81.9 mM Na2HPO4, 18.1 mM NaH2PO4) at 4L overnight. Samples were sequentially treated with Proteinase K (Merck), hydrochloric acid, and acetic acid. Subsequently, sections of the slides were dehydrated with 70%, 80%, 90%, and 100% ethanol and air-dried for at least 20 min. 900ng of RNA probe (**Table S2**) in 150 µl of Hybridization solution [10 mM Tris-HCl buffer, pH7.6 containing 50% formamide, 200 mg/ml tRNA, 1× DenhardtLs solution (Wako, Japan), 10% dextran sulfate, 600 mM NaCl, 0.25% sodium dodecyl sulfate, and 1 mM EDTA] was added to each slide and hybridized overnight at 50L. After washing with 2x saline sodium citrate (SSC) containing 50% formamide, sections were transferred into TNE buffer (10 mM Tris-HCl at pH 7.5, 0.5 M NaCl, 1 mM EDTA) and treated with RNase A (Sigma), followed by washing in TNE buffer, 2× SSC, and twice in 0.2×SSC. Blocking was performed with 1.5% blocking reagent (DIG Nucleic Acid kit) in DIG buffer1 (100mM Tris-HCl pH 7.5, 150mM NaCl) for 1 h. For fluorescent ISH, sections were treated with anti-DIG antibodies conjugated with peroxidase (Roche) diluted to 1:1000 in DIG buffer 1 for 30 min, followed by tyramide single amplification reaction solution [1/100 tyramide reagent (1 mg/ml succinimidyl ester, 10 mg/ml tyramine hydrochloride, 1% Triethylamine) in dimethyl formamide, 0.003% H_2_O_2_, 2% sodium dextran sulfate, 0.3 mg/ml 4-iodophenol] [69,70] for 15 min. For chromogenic ISH, sections were treated with anti-DIG antibodies conjugated with alkaline phosphatase (Roche) diluted 1:1000 in DIG buffer 1 for 30 min, followed by treatment with NBT/BCIP solution (DIG Nucleic Acid kit) diluted 1:50 in DIG buffer 3 (100mM NaCl, 200mM Tris-HCl pH9.5, 50mM MgCl_2_, 0.01% Tween20) for 3 h or overnight until signals were detected. EdU was detected in the samples injected with EdU. After blocking with 3% bovine serum albumin (BSA) in PBS, 200 μl of EdU detection buffer [Click-iT^TM^ EdU Cell Proliferation Kit (Invitrogen)] was added onto each slide and incubated for 30 min at room temperature. For all samples, after fixation in 4% PFA in PB and washing with PB, each slide was mounted in 70% glycerol. Images were captured using a light microscope BZ-X810 (Keyence).

### Enrichment of proliferating cells and non-proliferating cells by FACS

MBs of pupae were dissected in Dulbecco’s PBS (DPBS) and collected in a 1.5 ml tube containing DPBS (5 pupae per lot, 3 biological replicates). DPBS was replaced with 500 μl of enzyme solution (1 mg/ml papain, 1 mg/ml collagenase in DPBS) and stirred in a vortex for 5 min at 600 rpm at 37L. Cells were suspended by pipetting 200 times and centrifuged at 93×*g* for 10 min at 4L. After the supernatant was replaced with 500 μl of DPBS and the precipitate was resuspended, cell clumps and excess debris were removed by passing the suspension through a 20 μm cell strainer (PluriSelect). Then, 5 μl of 1 mg/ml Hoechst33342 (Invitrogen) in DPBS was added to the cell suspension, incubated at 25L for 20 min, and then centrifuged at 93×*g* for 10 min at 4L and replaced with 0.01% BSA/DPBS. 7-aminoactinomycin (7-AAD) was added to the solution at a final concentration of 1 μg/ml before sorting. SORPAria (BD Biosciences) was used to sort the cells according to the pattern of the forward/side scattered light signal (FSC/SSC), fluorescence signal of the dead cell staining reagent 7-AAD (fluorescent signal: perCP-Cy5-5), and nuclear staining reagent Hoechst 33342 (fluorescence signal: BUV395). First, cell fragments were removed based on cell size and particle size (intracellular complexity), followed by the removal of doublets based on the aspect ratio, and then dead cells. Finally, the fractions of the 2X and 4X nuclear phases were separated based on nuclear staining intensity. The sorted cells were collected by centrifugation at 93× g at 4L for 10 min and stored at −80L.

### RNA-seq analysis

Library synthesis and rRNA depletion were performed using the Takara SMARTer kit for Illumina sequencing, and sequences of approximately 20M reads × with 36bp pair end were obtained from each sample using NextSeq500 (Illumina). Trim-Galore [71] and hisat2 [72] were used to remove the adapter sequence from the reads and map them to the reference genome, respectively. StringTie [73] was used to obtain gene count data. Finally, the R (version 4.3.1) package edgeR [74] was used to search for genes with variable expression. Genes differentially expressed between the 2X and 4X fractions were identified as genes with a false discovery rate < 0.05, using the likelihood ratio test in a multifactor design with edgeR and over two-fold change in expression levels. A heat map was generated by calculating the ratio of reads per million (RPM) of the 2X fractions to that of the 4X fractions obtained from the same sample.

### GO enrichment analysis

The DEGs identified by RNA-seq were converted to one-to-one orthologs of *Drosophila* [56] and subjected to enrichment analysis using Metascape [75].

### Data accessibility

The raw RNA-seq data generated in the present study have been deposited in the DNA Data Bank of the Japan Sequence Read Archive database under the accession code PRJDB19871.

## Declaration of interests

The authors declare no competing interests.

## Acknowledgement

This work was supported by the Sasakawa Scientific Research Grant from The Japan Science Society, JST SPRING, Grant Number JPMJSP2108, and Japan Society for the Promotion of Science (JSPS) KAKENHI Grant Number 23K05875

## Supplemental information

FigureS1-3 and TableS1-2

---

**Figure S1 Time-lapse imaging of pupation in the artificial rearing condition**

Time-lapse images (10 min increment from left to right) of an individual from prepupa to pupa under artificial rearing conditions are shown. ‘−30 min’ indicates 30 min before pupation, and ‘+10 min’ indicates 10 min after pupation. The abdomen begins to move immediately before pupation and ceases to move after pupation.

**Figure S2 Identification of the pupal stage when each subtype is produced from neuroblasts**

Distribution of EdU signals in the brains of adult individuals injected with EdU at various pupal stages: 7 hap (A), 32 hap (B), 42 hap (C), 49 hap (D), 88 hap (E), 90 hap (F), and 108 hap (G). From left to right, EdU signals, *jhdk* FISH signals, merged signals, and schematic diagrams of EdU signals and each subtype are shown. Ca, calyx. Scale bar; 100 μm.

**Figure S3 Antisense probe-specific ISH signals of the genes suggested to be expressed in the proliferating cells or immature KCs.**

ISH results of *SoxN* (A), *optix* (B), *asense* (C), and *opa* (D) in the MBs of pupae with proliferating MB cells using antisense (left) and sense (right) probes. Black and grey arrows indicate strong and weak ISH signals, respectively, which were specifically detected using antisense probes. Ca, a calyx in the MB. Scale bar; 100 μm.

## Notes

### Competing Interest Statement

The authors have declared no competing interest.

