## Supplementary figures and images for "Analysis of molecular and cellular bases of honey bee mushroom body development"

### Supplementa Figure

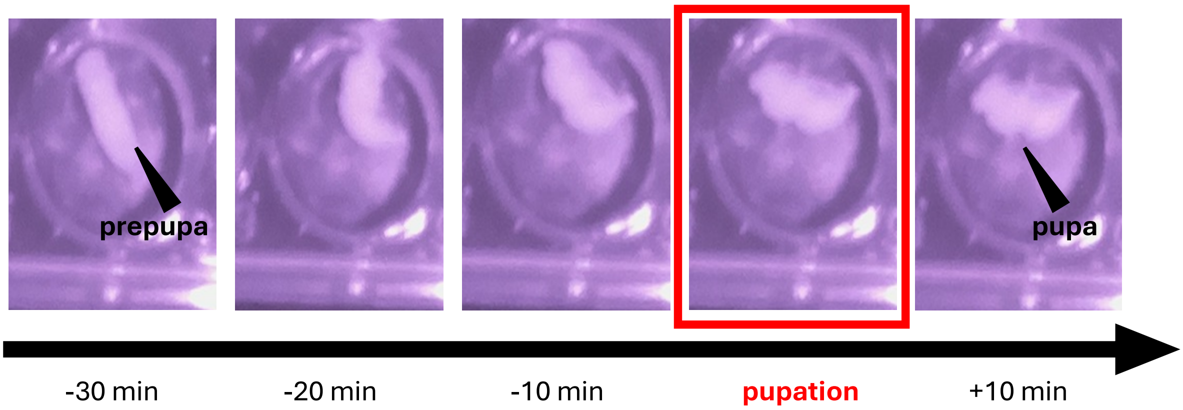


**FigureS1**

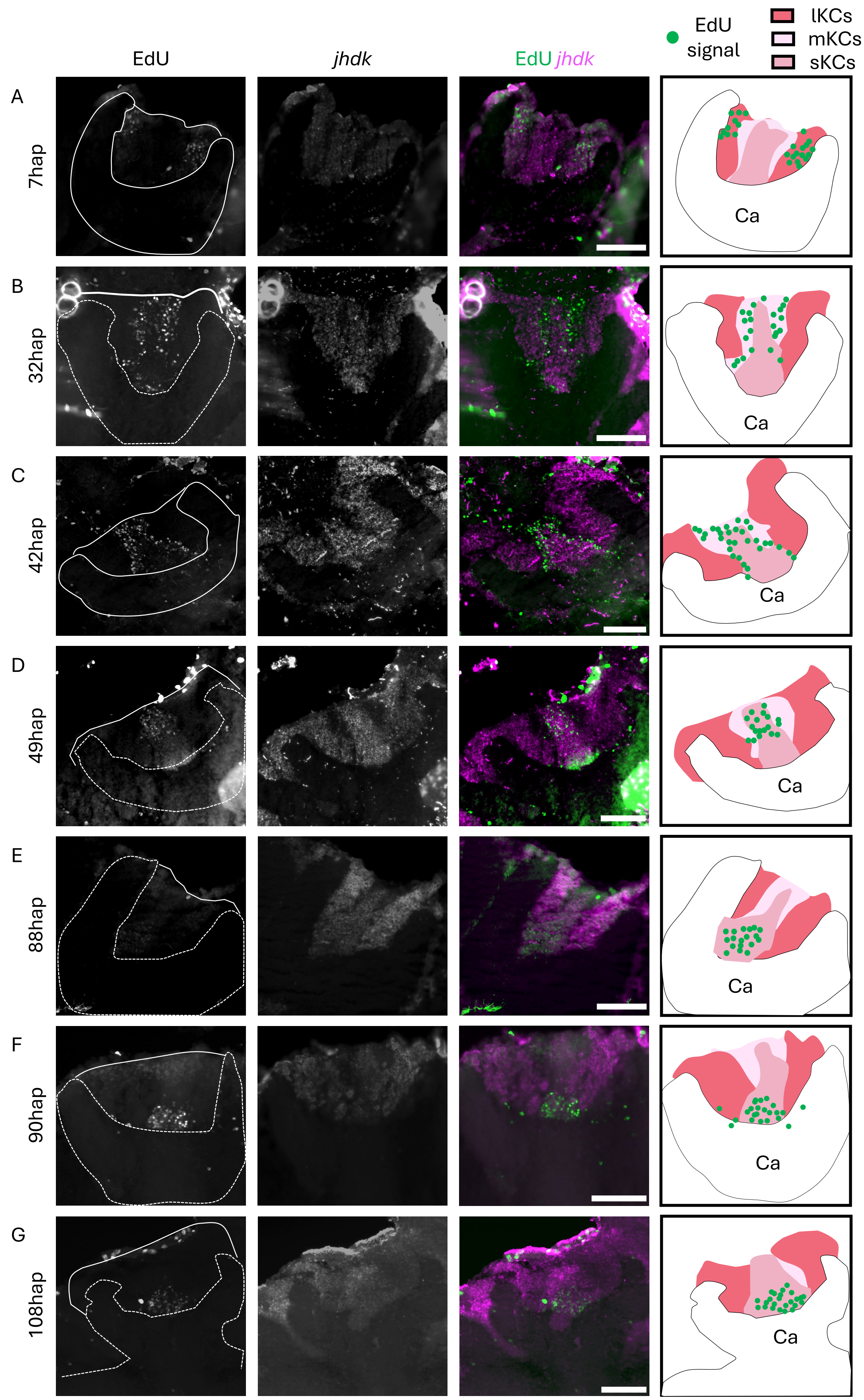

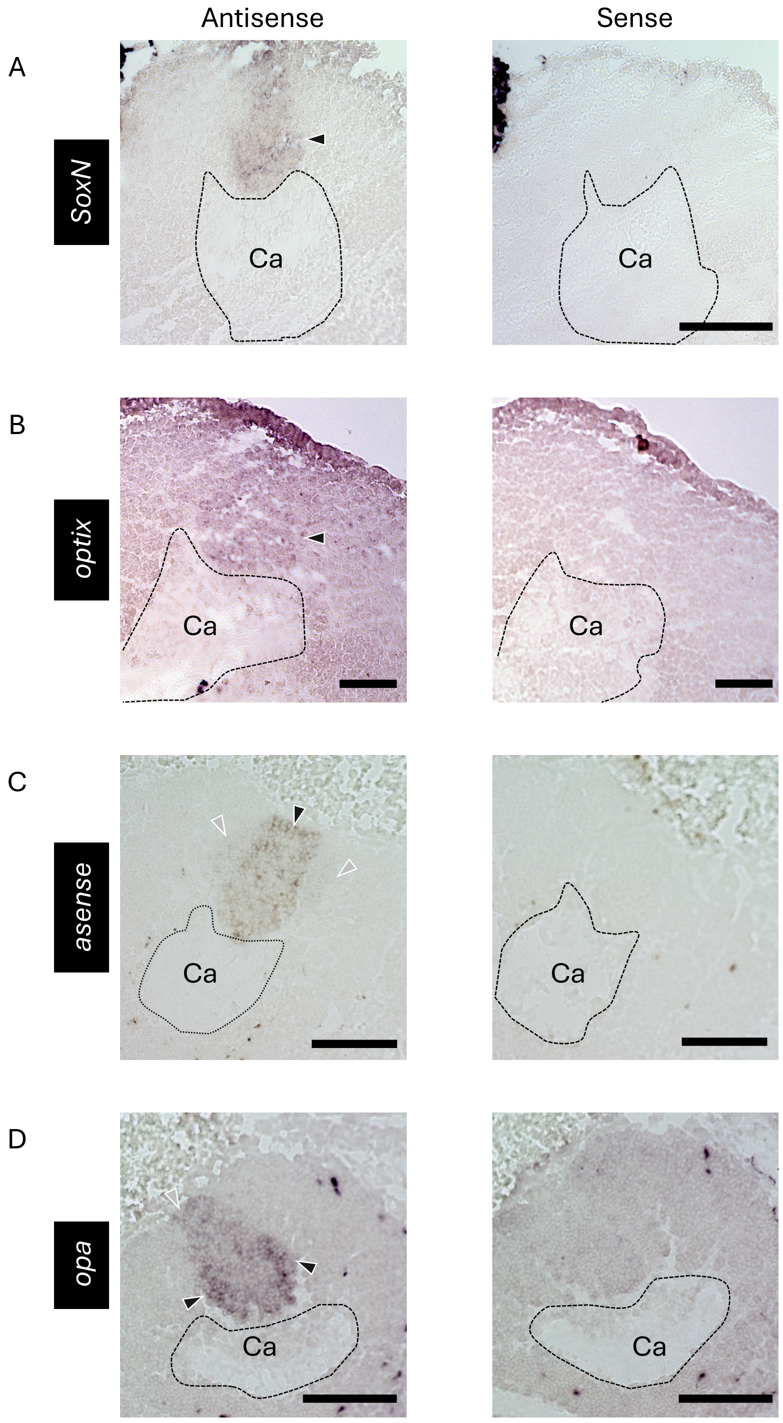
